# Basement membrane components define the microenvironment of aggregated fibroblasts in the skin and support their aggregation *in vitro*

**DOI:** 10.1101/2025.08.27.672757

**Authors:** Hiroki Machida, Jun Yokota, Hironobu Fujiwara

## Abstract

The dermal papilla (DP), a specialized fibroblast aggregate in mammalian skin, plays a pivotal role in hair follicle development and regeneration through epithelial‒mesenchymal interactions. While its aggregated configuration is critical for its function, the mechanism maintaining this organization has remained unclear. Here, we show that human DP cells are embedded in a basement membrane (BM)-like extracellular matrix (ECM) rather than in conventional interstitial fibrillar ECM, such as collagen I. This BM-like ECM occupies the intercellular space as a diffuse, mesh-like structure, with minimal cell‒cell adhesion. These fibroblasts interact with specific laminin isoforms containing α1, α2 or α4 chains via integrin α7β1, resulting in weak adhesiveness. *In vitro*, DP spheroids remained aggregated in a BM extract-based matrix, Matrigel, but dispersed in collagen I gel in an α1β1 integrin-dependent manner. Our findings suggest that the BM components within DP aggregates mediate cell‒ECM interactions and maintain the cohesive DP structure in the absence of strong cell‒cell contacts. These results reveal an unconventional role for the BM as an adhesive microenvironment that sustains fibroblast aggregation and offer a new perspective on mesenchymal tissue organization and *in vitro* culture substrates for DP cells.

## Introduction

Epithelial‒mesenchymal interactions are fundamental to the development and regeneration of various organs, including hair follicles, teeth, lungs, and kidneys [1–5]. These interactions occur within specialized microenvironments where extracellular matrix (ECM) components provide both structural support and biochemical cues essential for cell communication. In particular, the ECM at epithelial‒mesenchymal interfaces is thought to be highly specialized, contributing to tissue patterning and morphogenesis. However, the precise molecular composition and mechanisms by which the ECM regulates these interactions remain poorly understood.

The hair follicle serves as an excellent model for studying epithelial‒mesenchymal interactions because of its well-defined cellular populations and cyclic regenerative capacity [6–8]. A key player in this process is the dermal papilla (DP), a specialized aggregate of fibroblasts that orchestrate hair follicle development and regeneration. The DP forms during embryogenesis when fibroblasts aggregate in response to placode-derived morphogens, including FGF20 [9–13]. In mature skin, DP cells regulate the proliferation and differentiation of epithelial progenitors in the hair germ (HG) through various morphogens, such as Sostdc1, TGFβ2, FGF7, and Shh [14–18]. Although the DP undergoes dynamic remodeling throughout the hair cycle [19], its ability to retain an aggregated structure and appropriate cell number is crucial for maintaining proper regenerative capacity [20–22].

Given the importance of the DP in hair follicle development and regeneration, significant efforts have been made to establish effective *in vitro* DP cell culture methods, such as coculture with epithelial cells and supplementation with various growth factors [23, 24]. However, cultured DP cells often lose their ability to induce hair follicle generation, likely due to disruption of their native microenvironment and three-dimensional organization. Notably, restoring DP cell aggregation has been shown to recover their ability to induce hair follicle generation [25, 26]. To better recapitulate *in vivo* conditions, advanced coculture systems have been developed to mimic the mesenchymal condensation processes observed in feather buds, the dental papilla and the DP [27–31]. For example, 3D-printed collagen I gel microwells seeded with adult DP cells and neonatal epidermal keratinocytes have been used to generate hair follicle-like structures *in vitro*, albeit with low efficiency [32]. These findings underscore the importance of designing culture systems that incorporate both cell–cell and cell–ECM interactions occurring in their native *in vivo* environments to improve DP function.

Cell–ECM interactions, both *in vivo* and *in vitro*, have been implicated in DP formation and maintenance. The DP niche is thought to be composed of a specialized ECM that provides the biomechanical support and biochemical signals essential for DP function. While the precise molecular composition of the DP ECM has remained elusive for many years, recent systematic studies have begun to elucidate its unique features.

Although DP cells belong to the fibroblast lineage, they are encapsulated by basement membrane (BM)-like ECM components, including laminins and collagen IV [33–35]. This specialized ECM is thought to play a pivotal role in regulating DP cell behavior, including adhesion, aggregation, and signaling. Although integrin α9 has been identified as a DP-specific cell surface marker in mice [36], the specific molecular interactions between DP cells and their surrounding ECM and how these interactions contribute to DP function remain largely unexplored.

In this study, we investigated DP cell–ECM interactions, focusing on laminin isoforms as key regulators of DP cell adhesion and organization. Through tissue analyses and *in vitro* cell-based assays, we identified major laminin isoforms and their integrin-mediated interactions with DP cells. Three-dimensional culture assays of DP cell aggregates suggested that the BM-based ECM contributes to maintaining DP integrity as a compact aggregate. These findings increase our understanding of the specialized ECM in the DP niche and its role in cellular organization, providing fundamental insights into epithelial–mesenchymal interactions. Furthermore, such insights offer potential strategies for optimizing *in vitro* DP cell culture methods and advancing regenerative therapies.

## Results

### The microenvironment of human DP cells comprises BM components

Our previous report revealed that DP cells from mouse telogen hair follicles downregulate the expression of interstitial ECM-related genes and upregulate the expression of BM-related genes and are encapsulated by BM proteins [33]. To investigate whether similar ECM distribution patterns are present in the human DP, we first examined the tissue distribution of a major interstitial fibrillar ECM component, collagen I, and core BM molecules, perlecan and collagen IV, around human anagen scalp terminal hair follicles via immunohistochemical staining (Fig. 1A). In the dermis just beneath the epidermis, collagen I formed dense fiber bundles distributed in a reticular pattern (Fig. 1A, arrowheads in the upper dermis). In the deeper dermis, collagen I formed large bundles containing sparsely distributed collagen fibrils (Fig. 1A, middle dermis, lower dermis). In the dermal sheath surrounding the hair follicle, collagen I formed dense bundles parallel to the long axis of the hair follicle, and these bundles were distributed slightly away from the perlecan-positive BM (Fig. 1A, dermal sheath). In contrast, collagen I showed little deposition in the DP (Fig. 1A, hair bulb). Similarly, another major fibrillar interstitial ECM protein, fibronectin, was rarely distributed in the DP (Fig. S1A). Instead, perlecan and collagen IV were deposited around individual DP cells (Figs. 1A and 1B, hair bulb). These results suggest that although human DP cells are of the fibroblast lineage, they reside in BM components rather than typical interstitial ECM components.

**Fig. 1.**
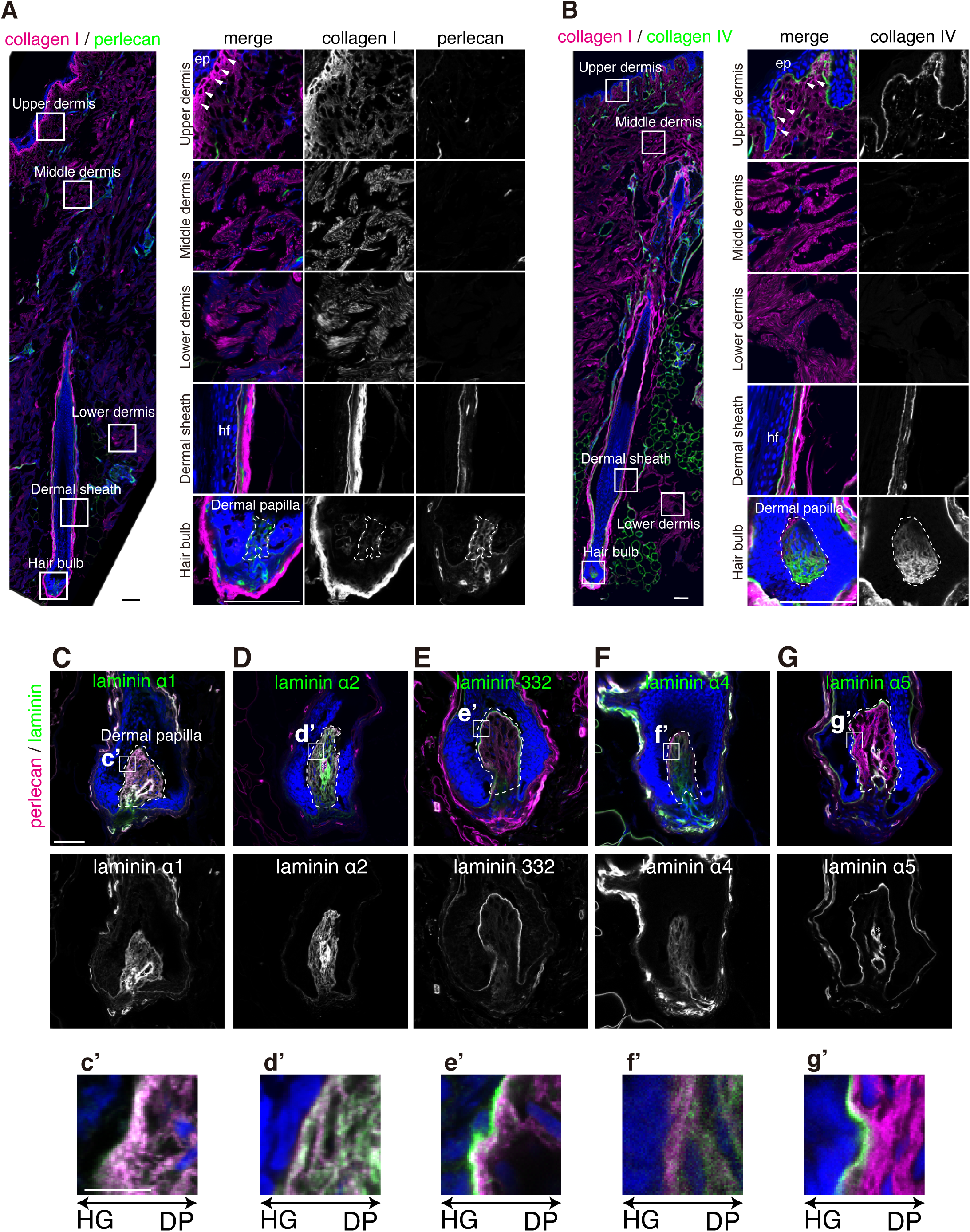
The tissue distribution of ECM proteins in human scalp hair follicles. (A and B) Immunohistochemical staining of collagen I (magenta) and perlecan (green in A) and collagen IV (green in B) in human female scalp skin with anagen terminal hair follicles. Nuclei were stained with DAPI (blue). The white squares indicate the region for the magnified images shown on the right. The arrowheads indicate dense fibrils of collagen I just beneath the epidermis, and the dashed lines indicate the DP. Scale bar, 100 µm. [ep, epidermis; hf, hair follicle]. (C–G) Immunohistochemical staining of perlecan (magenta) and laminin α chains (green) in the hair bulb region of human scalp anagen terminal hair follicles. The dashed lines mark the DPs. Asterisks indicate vasculature-like structures within the DP. Scale bar, 50 µm. Magnified images of the HG–DP interface are shown in the bottom panels (c’–g’). Scale bar, 10 µm.

Next, we investigated the localization of laminin α chains within the DPs of human terminal hair follicles. Laminin α chains are components of heterotrimeric laminin molecules, which mediate cell–BM interactions by binding to cell surface integrins and other receptors. Laminin α1, α2, and α4 chains were deposited in a mesh-like pattern surrounding individual DP cells (Figs. 1C, 1D, and 1F). They were also deposited at the tissue boundary between the hair germ and DP but were localized more toward the DP than the perlecan-positive BM (Figs. 1c’, 1d’ and 1f’). In contrast, laminin-332, which contains the laminin α3 chain, was deposited linearly at the hair germ (HG)–DP junction but was localized closer to the hair germ epithelium than the DP and was not detected within the DP (Figs. 1E and 1e’). The laminin α5 chain was deposited at the hair germ-DP junction and in the BM of blood vessel-like structures within the DP (Fig. 1G). At the HG-DP boundary, the laminin α5 chain was localized closer to the hair germ epithelium than the DP and was not detected within the DP (Fig. 1g’). These results suggest that DP cells interact with laminins containing α1, α2, and α4 chains, which form a mesh-like structure inside the DP.

In mouse telogen hair follicles, there are two BM-like structures: the mesh BM with the laminin α4 chain surrounding each DP cell and the hook BM with the laminin α5 chain protruding from the tissue boundary into the DP [33]. We confirmed that in human telogen facial vellus hair follicles, the laminin α5 chain similarly extended from the tissue boundary into the DP, whereas in anagen follicles, it was confined to the tissue boundary facing the hair follicle epithelium and was not observed on the dermal side (Fig. S1B).

In summary, these results suggest that human DP cells reside in a BM protein-rich environment, similar to that observed in mouse hair follicles.

### Potential cellular origins of laminins around the DP

To explore the cellular origin of laminins, we analyzed the gene expression of laminins using single-cell RNA sequencing (scRNA-seq) data from human abdominal skin containing hair follicles [37] (Fig. 2A). Among the laminins that were distributed around the DP, *LAMA1*, encoding laminin α1, was not detected in any cell population in the skin, including DP cells (FB-DP). *LAMA2*, encoding laminin α2, was expressed in FB-DP, but its highest expression was observed in lower dermal sheath fibroblasts (FB-LDS). Interestingly, the laminin α2 protein was only weakly detected around the lower dermal sheath but substantially accumulated within the DP (Fig. 1D), suggesting the presence of selective retention or recruitment mechanisms within the DP. Similarly, although *LAMA4*, encoding laminin α4, was highly expressed in Schwann cells (SCH), lymphatic endothelial cells (LEC), and broadly in fibroblast subpopulations, its expression was low in FB-DP and FB-LDS (Fig. 2A), suggesting a similar mechanism of selective localization within the DP. Telogen phase-specific laminin α5, encoded by *LAMA5*, was highly expressed in the basal outer root sheath cells (ORS) and vascular endothelial cells (VEC), but its expression was low in FB-DP and FB-LDS. These results suggest that laminins surrounding DP cells are derived primarily from dermal cells around the DP rather than from the DP cells themselves.

**Fig. 2.**
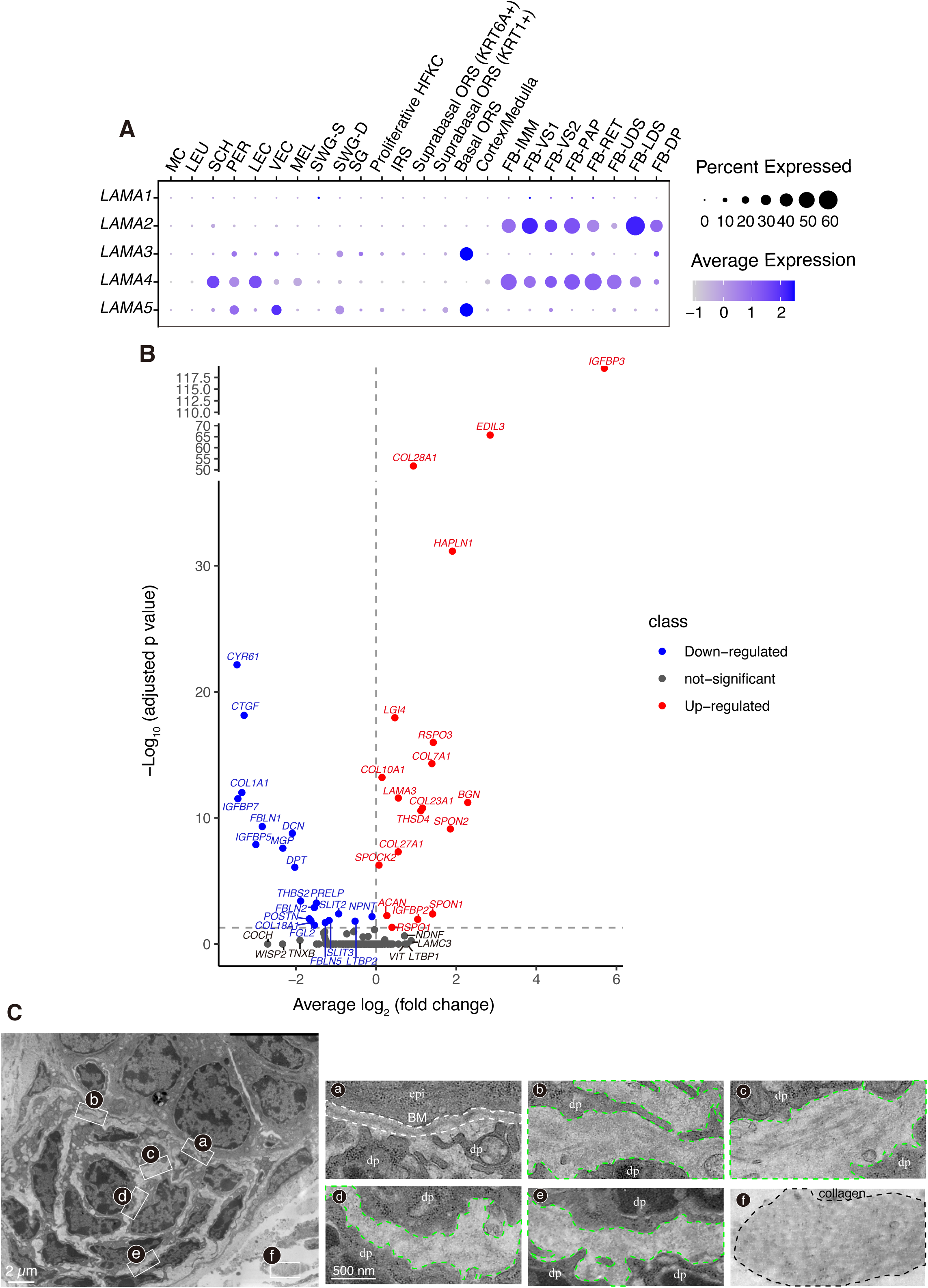
Distinct ECM gene expression and ultrastructural ECM organization in the DP. (A) Dot plot showing the gene expression of laminin α chains in the scRNA-seq data of human abdominal skin [37]. [MC, mast cells; LEU, leukocytes; SCH, Schwan cells; PER, pericytes; LEC, lymphatic endothelial cells; VEC, vascular endothelial cells; MEL, melanocytes; SWG-S, secretory cell comprising the sweat gland; SWG-D, duct cells comprising the sweat gland; SG, sebaceous gland; Proliferative HFKC, proliferative hair follicle keratinocyte; IRS, inner root sheath; ORS, outer root sheath; Suprabasal ORS (*KRT6A+*); KRT6A-positive suprabasal cells in outer root sheath; Suprabasal ORS (*KRT1+*), KRT1-positive suprabasal cells in outer root sheath; Basal ORS, basal cells in outer root sheath; Cortex/Medulla, hair cortex cells and medulla; FB-IMM, immune fibroblasts; FB-VS1, vasculature-related fibroblasts 1; FB-VS2, vasculature-related fibroblasts 2; FB-PAP, papillary fibroblasts; FB-RET, reticular fibroblasts; FB-UDS, upper dermal sheath fibroblasts; FB-LDS, lower dermal sheath fibroblasts; FB-DP, DP cells]. (B) Volcano plot showing upregulated and downregulated ECM genes in the DP cells (FB-DP) compared with other dermal fibroblasts, which constitute a pooled population of all dermal fibroblast subpopulations except FB-DP. Differential expression analysis was performed via model-based analysis of single-cell transcriptomics (MAST) [78] with the human homologs of the mouse core matrisome genes defined in Tsutsui et al. [33, 38]. The x-axis represents log_2_-fold change (FC) values of average expression (DP/other fibroblasts), and the y-axis corresponds to the log_10_ adjusted *p*-value derived from MAST with Bonferroni correction. The vertical gray dashed line indicates FC = 0, and the horizontal gray dashed line indicates the adjacent *p*-value threshold of 0.05. (C) Representative transmission electron microscopy (TEM) images of the DP in adult mouse telogen dorsal skin [33]. The white squares indicate the regions magnified in the images on the right. The white dashed line indicates the BM at the hair germ–DP interface; the green dashed lines denote extracellular spaces within the DP; and the black dashed line outlines collagen fibers. [epi, epidermis; dp, dermal papilla]

### Differentially expressed matrisome genes in human DP cells

To further explore the ECM composition of the DP, we analyzed human matrisome gene expression [33, 38] (Fig. 2B) and compared the average matrisome expression in FB-DP with that in all other skin fibroblast subtypes. Consistent with our protein localization analysis, *COL1A1*, which encodes the α1 chain of collagen I, was markedly lower in FB-DP, with a log_2_-fold change of -3.3, indicating approximately 10% of its expression in other fibroblast subtypes. FB-DP also had reduced expression of *DCN*, which encodes the collagen fiber formation-associated proteoglycan decorin [39]; *CTGF*, which encodes the fibrosis-related matricellular protein connective tissue growth factor [40]; and *FBLN1* and *FBLN2*, which encode fibrillins involved in microfibril formation [41]. In contrast, FB-DP exhibited increased expression of several nonfibril-forming collagen genes, including *COL7A1*, *COL10A1*, *COL23A1*, and *COL28A1* [42].

Many of the highly expressed matrisome genes were associated with tissue morphogenesis and growth factor signaling, such as *BGN*, which encodes the WNT-related proteoglycan biglycan [43]; *RSPO1*, *RSPO2*, and *RSPO3*, which encode the WNT agonists R-spondins that promote hair follicle growth [44, 45]; *SPON1* and *SPON2*, which encode F-spondins involved in morphogenesis through TGF-β activation [46]; and *IGFBP3*, which encodes insulin growth factor binding protein-3, a regulator of hair follicle epithelial proliferation and differentiation [47, 48]. These findings reveal that the expression of nonfibril-forming collagens and morphogenesis-related ECM proteins is upregulated, and the expression of fibrillar ECM components is downregulated in human DP cells.

### The ECM within the DP forms an amorphous ultrastructure

To further investigate the ultrastructure of the ECM surrounding DP cells, we reanalyzed electron microscopy images of mouse telogen hair follicles from a previous study (Fig. 2C) [33]. In the DP, although the cells were packed, direct cell‒cell contacts were rarely observed, and most of the extracellular space was filled with the ECM. A linear lamina densa structure was present at the tissue boundary between the hair germ and the DP (Fig. 2C, a), but it was absent within the DP (Fig. 2C, b–e). Compared with the other regions of the DP, the region of the DP closer to the tissue boundary with the hair follicle epithelium (Fig. 2C, b and c) presented a more extensive distribution of electron-dense structures (Fig. 2C, d and e). In contrast, the remaining extracellular regions of the DP displayed a relatively scattered distribution of coarse fibrillar or electron-lucent amorphous structures (Fig. 2C, d and e), which were distinct from typical collagen fibers or the lamina densa structure (Fig. 2C, f). These observations suggest that, despite being rich in BM components, the ECM of the DP does not form the typical BM ultrastructure.

### DP cells weakly adhere to laminins containing α1, α2, or α4 chains

To determine whether DP cells adhere to laminins surrounding DP cells, solid-phase cell adhesion assays were performed using purified recombinant laminin proteins and cultured human scalp-derived DP cells. We used five types of full-length and E8 fragments of human laminin proteins that differ in the type of α chain responsible for cell adhesion activity (laminin-111, -111 E8, -211, -221 E8, -332, -332 E8, -411, -411 E8, -511 and -511 E8), as well as the major interstitial fibrillar ECM proteins fibronectin and collagen I [49]. We measured the number of adhered cells after 30 minutes of incubation in culture dishes coated with these ECM proteins at different concentrations (Fig. 3A). To compare the cell adhesion activity of the ECM proteins, we calculated the concentration of ECM proteins required to induce 50% of the peak number of adherent cells (ED50) from the dose‒response curves (Fig. 3B, Table 1). Lower ED50 values indicate greater adhesiveness.

**Fig. 3.**
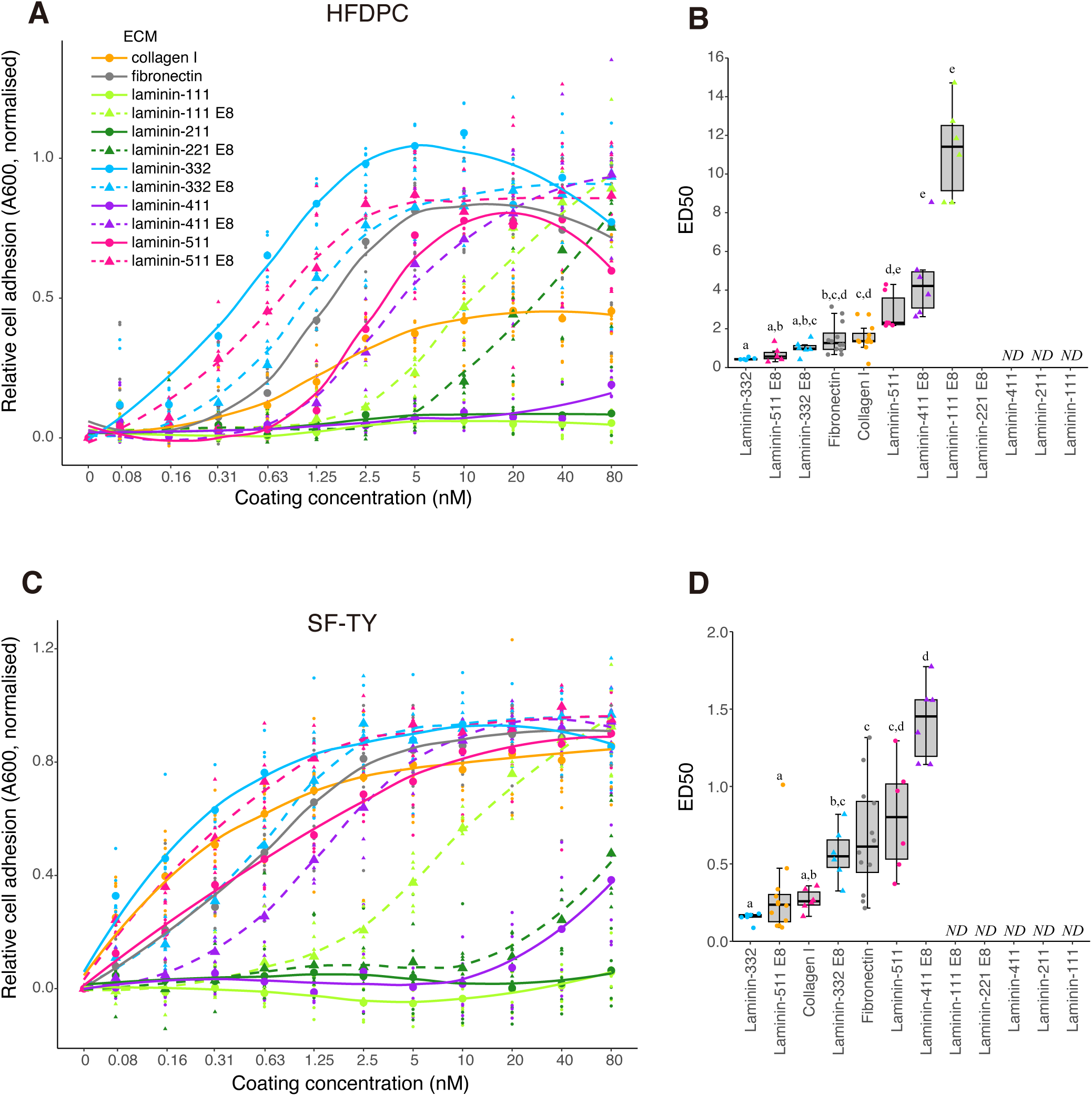
Adhesion of DP cells and skin fibroblasts to ECM proteins. (A and C) Dose‒response curves showing the absorbance (A600) reflecting the relative number of adherent cells in solid-phase cell adhesion assays performed with HFDPCs (A) or the human normal skin fibroblast cell line SF-TY (C). The smaller dots represent the normalized values of the number of adhered cells, which were calculated relative to the maximum effect observed on fibronectin for each technical replicate. Larger dots indicate the mean values of the number of adhered cells on each ECM protein, and the colored lines are their fitting curves. (A and C) *n* = 12 wells pooled from 4 independent experiments for collagen I and fibronectin; *n* = 6 wells pooled from 2 independent experiments for full-length laminins and laminin E8 fragments. (B and D) Box plots representing the median effective dose 50 (ED50) of cell adhesive activities of each ECM protein against HFDPCs (B) and SF-TY cells (D). The box shows the interquartile range with the median; whiskers extend to the 1.5 interquartile range. Statistical comparisons were made using the Kruskal‒Wallis test followed by Dunn’s post hoc test. Adjusted *p*-values were calculated via the Benjamini‒Hochberg method. Groups with different letters (a–e) indicate statistically significant differences in ED50 values (*p*<0.05). ND: the ED50 could not be determined due to the absence of a plateau.

**Table 1.**
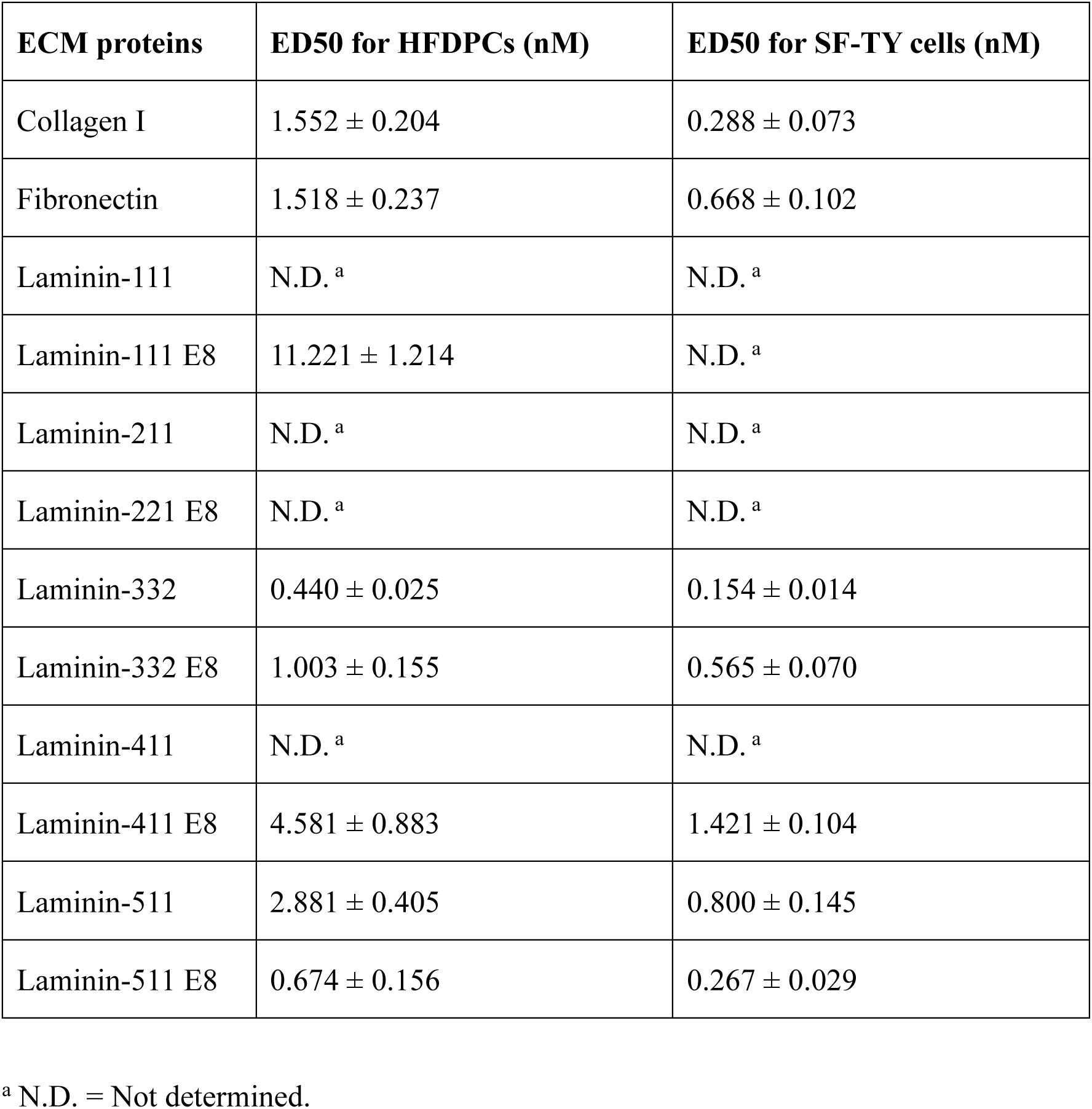
ED50 values of the cell adhesive activities of various ECM components.

| ECM proteins | ED50 for HFDPCs (nM) | ED50 for SF-TY cells (nM) |
| --- | --- | --- |
| Collagen I | 1.552 ± 0.204 | 0.288 ± 0.073 |
| Fibronectin | 1.518 ± 0.237 | 0.668 ± 0.102 |
| Laminin-111 | N.D.<sup>a</sup> | N.D.<sup>a</sup> |
| Laminin-111 E8 | 11.221 ± 1.214 | N.D.<sup>a</sup> |
| Laminin-211 | N.D.<sup>a</sup> | N.D.<sup>a</sup> |
| Laminin-221 E8 | N.D.<sup>a</sup> | N.D.<sup>a</sup> |
| Laminin-332 | 0.440 ± 0.025 | 0.154 ± 0.014 |
| Laminin-332 E8 | 1.003 ± 0.155 | 0.565 ± 0.070 |
| Laminin-411 | N.D.<sup>a</sup> | N.D.<sup>a</sup> |
| Laminin-411 E8 | 4.581 ± 0.883 | 1.421 ± 0.104 |
| Laminin-511 | 2.881 ± 0.405 | 0.800 ± 0.145 |
| Laminin-511 E8 | 0.674 ± 0.156 | 0.267 ± 0.029 |

<sup>a</sup> N.D. = Not determined.

With respect to DP cell adhesion to the interstitial fibrillar ECM, the ED50 was 1.6 nM for collagen I and 1.5 nM for fibronectin (Fig. 3B). The maximum number of DP cells adhering to collagen I was approximately half that observed with fibronectin (Fig. 3A). With full-length laminins, DP cells adhered to laminins containing the α3 chain (laminin-332) and α5 chain (laminin-511), with ED50 values of 0.4 nM and 2.9 nM, respectively. Laminin-332 exhibited the highest degree of cell adhesion-inducing activity. In contrast, DP cells did not adhere to laminins containing the α1 chain (laminin-111), α2 chain (laminin-211), or α4 chain (laminin-411) within the range of coating concentrations used in this study. With laminin E8 fragments, DP cells adhered to laminin-111 E8, laminin-221 E8, and laminin-411 E8, although the cell adhesive activities were lower than those of other full-length and E8 laminins containing α3 and α5 chains and fibronectin. The ED50 values for laminin-411 E8 and laminin-111 E8 were 4.6 nM and 11.2 nM, respectively. DP cells begin to adhere to laminin-221 E8 at a coating concentration of 5 nM. However, the number of adherent cells did not reach its maximum even at concentrations as high as 80 nM. When these results are considered alongside the tissue distribution data (Figs. 1C–1G), they suggest that DP cells weakly adhere to laminins containing α1, α2, and α4 chains, which form a mesh-like structure surrounding each DP cell. Moreover, the results suggest that DP cells strongly adhere to laminins containing the α5 chain, which appears in the hook-BM of telogen hair follicles.

To further investigate how the cell–ECM adhesion characteristics of DP cells differ from those of other skin fibroblasts, we conducted the same cell adhesion assays using a normal human skin fibroblast cell line, SF-TY (Fig. 3C). Unlike DP cells, SF-TY cells adhered well to collagen I, with the maximum number of adherent cells comparable to that observed with fibronectin and an ED50 of 0.3 nM (Figs. 3C and 3D). The ED50 for fibronectin was 0.7 nM, which was also lower than that of DP cells. SF-TY also adhered strongly to laminin-332 and laminin-511, with ED50 values of 0.2 nM and 0.8 nM, respectively (Fig. 3D). Furthermore, SF-TY adhered to laminin-411, to which DP cells showed no adhesion, starting at a coating concentration of 10 nM. However, the number of adherent cells did not reach a plateau within the concentration range of 10–80 nM (Fig. 3C). In the adhesion assays with laminin E8, SF-TY adhered to laminin-332 E8, -411 E8, and -511 E8, with ED50 values of 0.6 nM, 1.4 nM, and 0.3 nM, respectively (Figs. 3C and 3D). SF-TY also adhered to laminin-111 E8 and -221 E8 starting at 1.25 nM and 10 nM, respectively. However, it did not reach a plateau, and we could not calculate ED50 values. The adhesion of SF-TY to laminin-221 E8 was weaker than that of DP cells, and even at a coating concentration of 640 nM, it did not reach a plateau (Fig. S2).

Taken together, these findings demonstrate that DP cells exhibit distinct ECM adhesion properties compared with those of other skin fibroblasts. DP cells showed weak adhesion to laminins containing the α1, α2 and α4 chains, which form a mesh-like structure surrounding each DP cell, while strongly adhering to laminins containing the α3 and α5 chains, which are not localized around the DP cells. In contrast, normal skin fibroblasts adhered well to both collagen I and laminins.

### Integrin α7 primarily mediates DP–laminin adhesion

Compared to SF-TY cells, DP cells exhibited different ECM adhesion properties. We thus hypothesized that these differences were due to variations in cell surface ECM receptors. To test this hypothesis, we analyzed integrin expression in DP cells and SF-TY cells via flow cytometry (Fig. 4A). DP cells presented increased levels of integrin α1 (collagen receptor) and α3 and α7 (laminin receptor) but decreased levels of integrin α2 (collagen receptor) and α6 (laminin receptor). To further investigate the expression patterns of ECM receptors in different skin fibroblast subpopulations, we analyzed scRNA-seq data obtained from human abdominal skin samples [37]. Among all the human fibroblast subpopulations classified by Yokota et al., FB-DP highly expressed *ITGA1* (integrin α1), *ITGA7* (integrin α7), *ITGA8* (integrin α8; RGD receptor), *ITGA9* (integrin α9; RGD-independent fibronectin receptor), *ITGAV* and *ITGB3* (integrin αVβ3; RGD receptor), *DAG1* (dystroglycan), and *SDC1*, *2* and *4* (syndecan) (Fig. 4B). In contrast, two major fibroblast subpopulations, reticular dermis fibroblasts (FB-RET) and papillary dermis fibroblasts (FB-PAP), presented lower levels of *ITGA1* but higher levels of *ITGA2* (integrin α2) (Fig. 4B). Immunohistochemical staining confirmed that integrins β1, α1, α3, α5, α7 α8, α9, and αV were distributed in the human DPs of scalp terminal hair follicles (Fig. 4C).

**Fig. 4.**
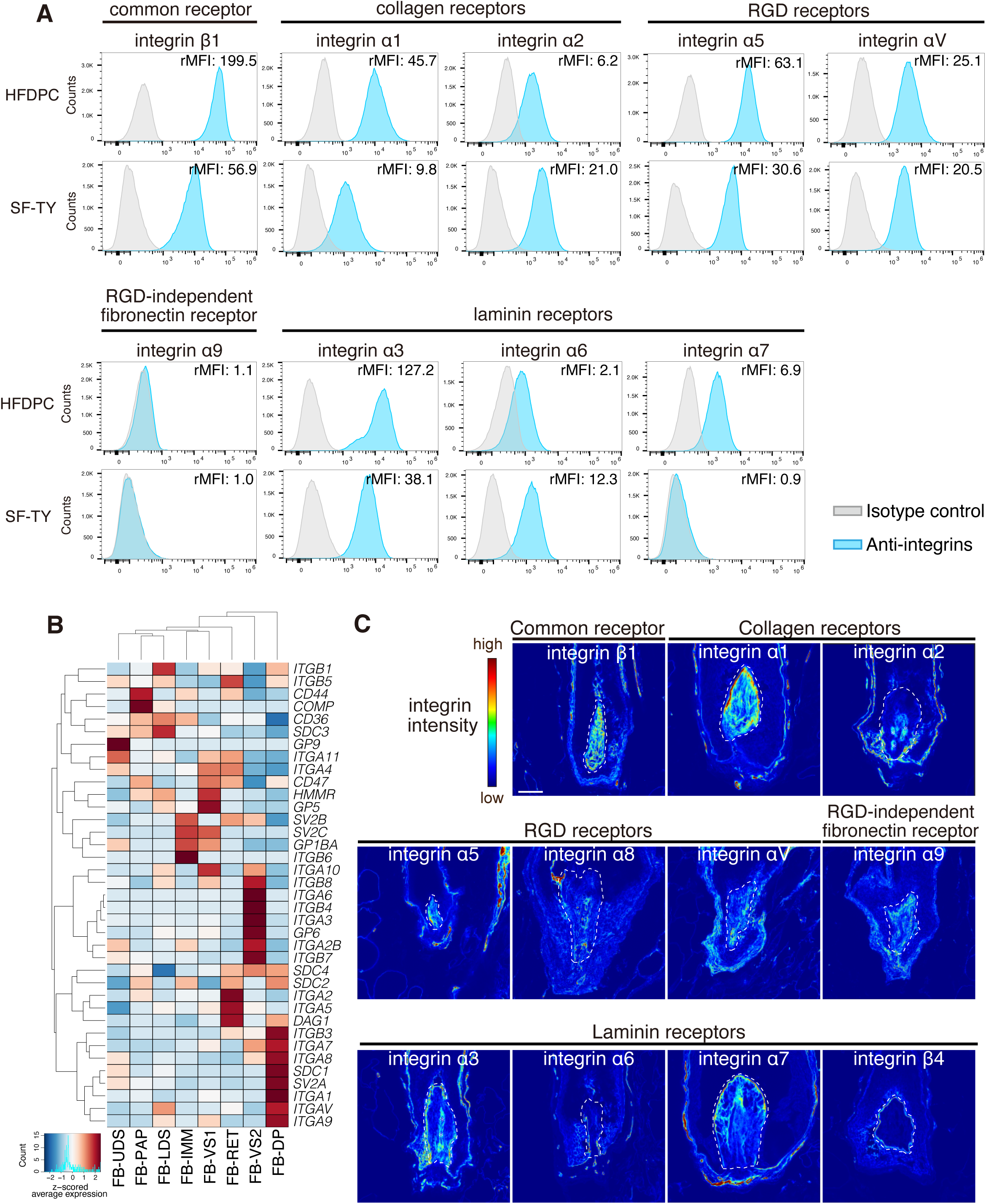
Differential expression of ECM receptors between DP cells and a skin fibroblast cell line. (A) Flow cytometry histograms showing integrin levels on the surface of HFDPCs and SF-TY cells. Relative-mean fluorescent intensities (rMFIs) normalized against the isotype control are shown at the top right of each histogram. Integrins are classified on the basis of their major ligands [52]. (B) Heatmap illustrating the hierarchical clustering of ECM receptor gene expression (KEGG: hsa04512) in human skin fibroblasts. scRNA-seq data from human abdominal skin [37] were analyzed. Z-scored average gene expression values for each fibroblast subpopulation are shown. [FB-PAP, papillary fibroblast; FB-RET, reticular fibroblast; FB-UDS, upper dermal sheath; FB-LDS, lower dermal sheath; FB-IMM, immune fibroblast; FB-VS1, vasculature-related fibroblasts 1; FB-VS2, vasculature-related fibroblasts 2; FB-DP, DP cells] (C) Representative immunohistochemical images showing integrin localization in the hair bulb region of human scalp terminal hair follicles. Integrins are classified on the basis of their major ligands [52]. The intensity of each integrin is indicated using a heat color scale (top left). The dashed lines indicate the DP. Asterisks indicate vasculature-like structures within the DP. Scale bar, 50 µm.

Next, to identify integrins that mediate DP cell adhesion to various ECM proteins, we performed cell adhesion inhibition assays using integrin-blocking antibodies. The adhesion of both DP cells and SF-TY cells to fibronectin was inhibited by antibodies against integrin α5 and β1 (Fig. 5A). The adhesion of DP cells to collagen I was inhibited by antibodies against integrins α1 and β1, whereas that of SF-TY cells was inhibited by antibodies against integrins α2 and β1 (Fig. 5B). These results indicate that DP cells use different types of integrins than general fibroblasts do for adhesion to collagen I. Both integrin α1β1 and α2β1 are known to bind to collagen types I and IV. However, integrin α1β1 has a greater affinity for collagen IV, whereas integrin α2β1 has a greater affinity for collagen I [50, 51]. Since collagen IV is known to be one of the major components of the BM and surrounds DP cells (Fig. 1B), this finding suggests that the mode of collagen adhesion in DP cells involves BM-type collagen rather than interstitial collagen.

**Fig. 5.**
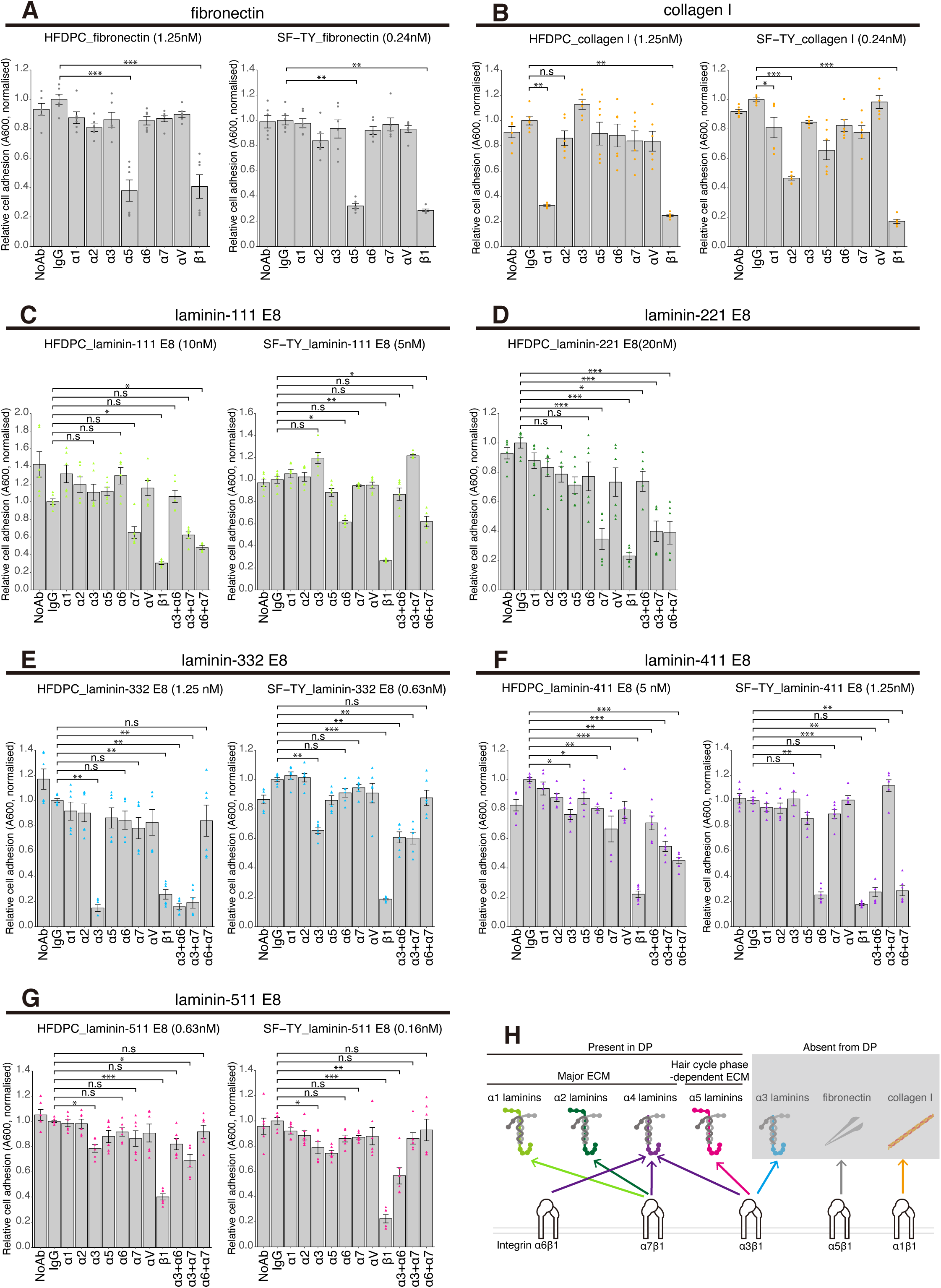
Identification of integrins mediating DP-ECM adhesion. (A–G) Bar graphs showing the results of cell adhesion inhibition assays for fibronectin (A), collagen I (B), laminin-111 E8 (C), laminin-221 E8 (D), laminin-332 E8 (E), laminin-411 E8 (F), and laminin-511 E8 (G). Each ECM protein was coated at the approximate ED50 coating concentrations. The absorbance at 600 nm reflects the relative number of adhered cells, normalized to the no-antibody (No Ab) control within each technical replicate. The data are presented as the means ± standard deviations (SDs). Statistical comparisons were performed via the Kruskal‒Wallis test followed by Dunn’s post hoc test. Adjusted *p*-values were calculated via the Benjamini‒Hochberg method: \**p*<0.05, \*\**p*<0.01, \*\*\**p*<0.001. *n* = 6 wells pooled from 2 independent experiments. (H) Schematic summary of how ECM protein localization and integrins mediate DP-ECM adhesion.

Given that cell adhesion to laminins is mediated primarily by integrins containing the α3, α6, and α7 subunits, we examined the effects of blocking antibodies against these integrin subunits [52]. The adhesion of DP cells to both laminin-111 E8 and laminin-221 E8 was inhibited by antibodies against integrin α7 and β1 (Figs. 5C and 5D). The adhesion of DP cells to laminin-332 E8 was inhibited by antibodies against integrin α3 and β1 (Fig. 5E). DP cell adhesion to laminin-411 E8 was inhibited weakly by antibodies against the α3, α6, and α7 subunits individually but was more strongly inhibited when any two of these integrins were simultaneously blocked—particularly when the antibody against α7 was included (Fig. 5F). Adhesion to laminin-511 E8 was inhibited by antibodies against integrin α3 and β1 (Fig. 5G). The results of the SF-TY cell experiments were slightly different. The inhibitory effect of integrin α6 on adhesion to laminin-111 E8, laminin-411 E8, and laminin-511 E8 was observed (Fig. 5C–G). Taken together, these results suggest that the adhesion of DP cells to laminins is mediated primarily by integrins α3β1 and α7β1, whereas the adhesion of SF-TY cells to laminins is mediated by integrins α3β1 and α6β1 (Fig. 5H). Notably, the adhesion of DP cells to laminin isoforms containing α1, α2, and α4 chains, which are distributed in the DP in the tissue, is mediated mainly by the DP-specific laminin-binding integrin α7 (Figs. 1B, 1C, 1E, 4A, 4B, 5C, 5D, and 5F).

### BM components can maintain the compact aggregate structure of the DP

In recent studies, collagen I gel is often used as a substrate for DP cultures [29, 31, 32]. However, our investigation of the molecular basis of cell–ECM interactions suggests that DPs are largely devoid of interstitial ECM, indicating that collagen I-based substrates may not fully recapitulate the native DP niche. Given that each DP cell is instead encapsulated by BM components, we investigated the biological significance of this specialized ECM environment by embedding DP spheroids in collagen I gel and Matrigel, the latter of which provides a biochemical environment similar to that of native BM (Fig. 6A).

**Fig. 6.**
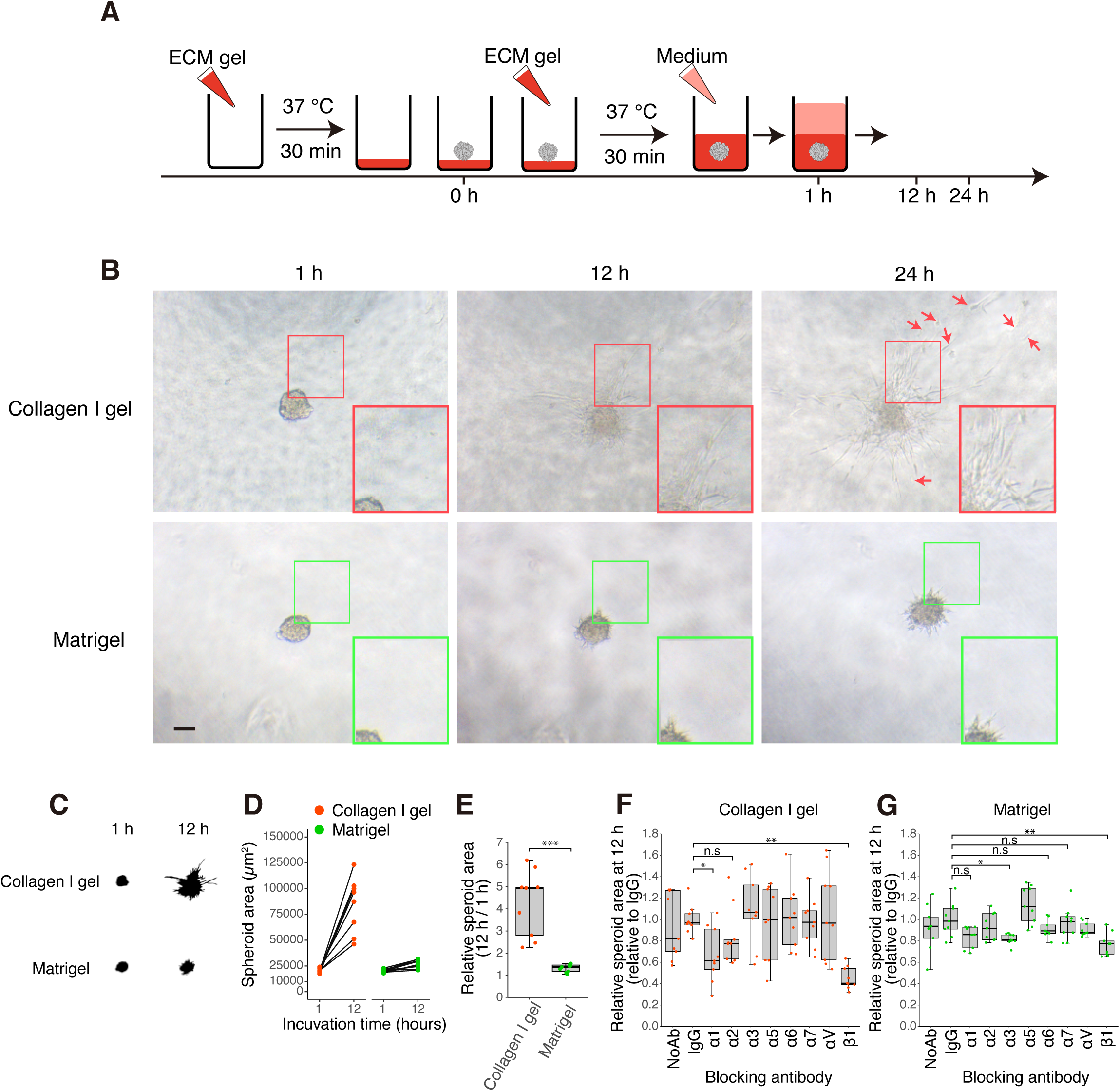
Matrigel maintains DP spheroid integrity. (A) Schematic of the spheroid invasion assay. DP spheroids, each composed of 300 HFDPCs, were placed on an ECM gel layer and overlaid with an additional ECM gel layer. The spheroids were cultured and observed every 12 hours. (B) Phase-contrast images of DP spheroids cultured in collagen I gel (top) and Matrigel (bottom). The colored squares indicate regions magnified in the bottom right images. Scale bar: 100 µm. (C) Binary-masked images highlighting the spheroid area. (D) Line graphs showing the changes in spheroid area over time in collagen I gel (red) and Matrigel (green). (E) Box plots showing the relative spheroid area at 12 h, normalized to the initial time point (1 h). The box shows the interquartile range with the median; whiskers extend to the 1.5 interquartile range. Two-tailed unpaired *t*-tests were used: \**p*<0.05, \*\**p*<0.01, and \*\*\**p*<0.001. *n* = 9 wells. (F and G) Box plots showing the relative spheroid area with integrin blocking antibodies in collagen I gel (F) and Matrigel (G). The box shows the interquartile range with the median; whiskers extend to the 1.5 interquartile range. Statistical comparisons were performed using the Kruskal‒Wallis test followed by Dunn’s post hoc test. Adjusted *p*-values from the Benjamini‒Hochberg method are shown: \**p*<0.05, \*\**p*<0.01, \*\*\**p*<0.001. *n* = 9 wells pooled from 3 independent experiments.

Matrigel is composed mainly of laminins (predominantly laminin-111 with relatively small amounts of laminin α3, α4, α5, and β2 chains), collagen IV, and perlecan [53, 54]. When embedded in collagen I gel, the DP cells began to invade the gel within 12 hours, and by 24 hours, the cells at the spheroid periphery had dispersed as individual cells (Figs. 6B–E). In contrast, although DP spheroids embedded in Matrigel extended short protrusions, they remained compact (Figs. 6B–E). These findings suggest that BM molecules contribute to maintaining DPs as compact aggregates.

To determine whether the integrins that mediate DP cell–ECM adhesion also regulate spheroid dispersion, we treated DP spheroids with integrin-blocking antibodies. In the collagen I gel, invasion was inhibited by antibodies against integrins α1 and β1 (Fig. 6F), indicating that adhesion to collagen I via integrin α1β1 promotes DP spheroid dispersion. In Matrigel, none of the tested antibodies—including those targeting the DP-specific laminin-binding integrin α7β1—induced spheroid invasion, suggesting that no integrins actively maintain DP spheroids in compact aggregates (Fig. 6G). Instead, blocking integrin α3 and β1 tended to inhibit protrusion formation (Fig. 6G).

Taken together, our findings suggest that BM molecules in the DP provide a structural scaffold that maintains the DP in an aggregated form. While sufficient integrin-mediated adhesion occurs in both environments, Matrigel may limit the transmission of traction forces necessary for DP spheroid dispersion [55], reinforcing its role in sustaining DP aggregation.

## Discussion

In this study, we revealed key features of human DP-ECM interactions (Fig. 7): 1) the ECM surrounding DP cells is enriched in BM-type ECM molecules and largely devoid of fibrillar interstitial ECM; 2) specific integrins mediate interactions between DP cells and these BM components; and 3) Matrigel helps maintain DP spheroids as compact aggregates, whereas collagen I gel promotes their dispersion. Given that mesenchymal fibroblast aggregation is essential for the development and regeneration of various organs, including hair follicles, feathers, and the intestine [11, 27, 56, 57], our findings suggest that integrin-mediated cell–ECM interactions may play a critical role in pattern formation.

**Fig. 7.**
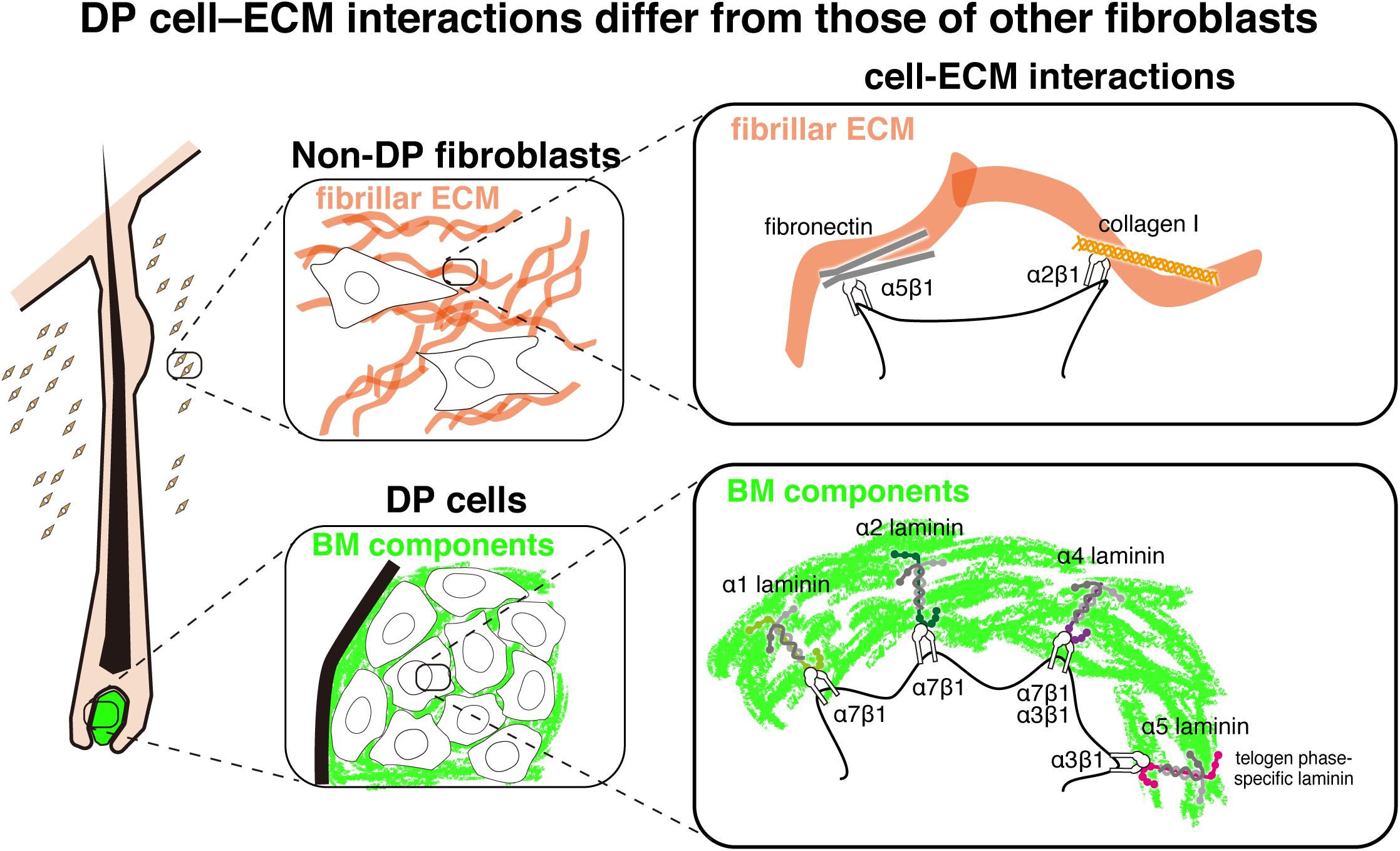
Graphical summary of DP–ECM interactions. This illustration highlights the spatial distribution of ECM molecules and their interactions with cells in the human dermis. Non-DP fibroblasts are typically surrounded by interstitial fibrillar ECM components, including fibronectin and collagen I, which interact primarily with integrins α5β1 and α2β1, respectively. In contrast, the DP microenvironment is characterized by a reduced presence of interstitial fibrillar ECM and an enrichment of BM components. Laminins containing α1, α2, and α4 chains appear to serve as the major adhesive substrates for DP cells, although these interactions—mediated by integrins α3β1 and α7β1—are weak. Laminin α5 has a unique hook-like distribution in the DP during the resting phase of the hair regeneration cycle in a phase-dependent manner. Additionally, it shows stronger adhesiveness to DP cells than other laminins, primarily via integrin α3β1.

### Characteristics of the ECM within the DP

The ECM of the DP is clearly distinct from that of the surrounding dermis. While the dermal ECM is primarily composed of interstitial fibrillar ECM molecules, such as collagen I and fibronectin, the DP-associated ECM is enriched with BM components, including laminins and collagen IV. Electron microscopy further revealed that DP cells are not tightly connected by cell‒cell adhesions but instead are separated by extracellular spaces filled with BM-like ECM. These findings suggest that DP aggregation relies more on cell–ECM interactions than on direct cell–cell contacts.

Unlike typical dense, sheet-like structures with well-defined lamina densa, the ECM surrounding DP cells appears more diffuse and mesh-like (Fig. 2C). This is potentially due to the presence of laminin α4, which lacks the LN domain required for stable laminin network assembly [58–60] and shows low integrin binding affinity [61, 62]. High coating concentrations of laminins containing α1, α2, or α4 are required to induce DP cell adhesion, suggesting that these weak interactions hinder the formation of a rigid laminin network. Given that collagen IV assembly depends on a preformed laminin network [60, 63, 64], it is plausible that such weak interactions contribute to a less polymerized, softer ECM environment in the DP. This softer matrix may in turn permit greater flexibility, which could support the dynamic morphological changes observed in the DP during the hair cycle, from spherical aggregates in telogen to elongated forms in anagen [8, 19, 65, 66].

### BM components maintain DP cells in an aggregated form

This potentially soft and fluidic BM-rich ECM may also facilitate the long-term aggregation of DP cells. *In vitro*, DP spheroids embedded in Matrigel—a soft, BM-rich matrix—remained compact despite extending short protrusions, whereas those in collagen I gels dispersed. Matrigel likely provides both a mechanical environment that limits traction forces and a weak but sufficient adhesive context through laminins. This *in vitro* behavior is consistent with *in vivo* ultrastructural features: DP cells show sparse cell‒cell adhesion, and their intercellular space is filled with ECM. The observation that spheroids resist dispersion in Matrigel, despite lacking strong cell‒cell junctions, raises the possibility that ECM components in the matrix infiltrate the spheroid and occupy the intercellular space, thereby contributing to cohesion. Together, these observations suggest that the ECM, rather than direct cell‒cell adhesion, functions as the main cohesive force maintaining DP aggregation. This concept aligns with the wetting model proposed by Huycke et al. 2024 [56], where the balance between surface tension and adhesion determines fibroblast aggregate stability. Consistent with this idea, Yang et al. 2023 [57] demonstrated that in avian dermal follicle condensate formation, increasing the stiffness of dermal cell clusters suppressed aggregate formation *in vitro*, further supporting the role of mechanical properties in DP cohesion.

### Integrins mediating DP–ECM adhesion

Our study identified integrin α7 as a major mediator of DP cell adhesion to BM-like ECM, specifically laminins containing α1, α2, and α4 chains. Ligand-binding assays with purified integrins and laminins indicate that integrin α7 has increased affinity for these laminins [61]. Its strong expression in DP cells suggests that it plays a significant role in maintaining DP integrity and regenerative function.

Integrin α7 is known for its role in skeletal muscles, where the BM composition changes from α2, α4, and α5 laminins during development to α2 laminins in mature tissue [67]. The integrin‒laminin interactions in embryonic skeletal muscle mirror those found in the DP, suggesting a conserved function across tissues. DP and skeletal muscle both originate from the mesoderm and depend on BM-mediated adhesion, despite diverging into distinct lineages. In skeletal muscle, integrin α7 knockout causes dystrophic phenotypes [68, 69]. In contrast, integrin α7 knockout or β1 integrin deletion in the dermis has not produced notable skin phenotypes [70, 71]. However, β1 integrin deletion via Col1a2-CreER reduces dermal thickness and collagen production in adult skin [72], suggesting context-specific roles in dermal compartments. Given the low collagen I expression of DP cells (Fig. 1 and 2B), β1 integrin may be retained in DP cells even in these knockout models. Thus, while the function of β1 integrin in DP cells *in vivo* remains unclear, its role likely differs from that in surrounding fibroblast populations.

In addition to laminin-mediated adhesion, our results indicate that DP cells also optimally adhere to BM-specific collagen, particularly collagen IV, through integrin α1β1. In contrast, SF-TY cells use integrin α2β1, which binds more strongly to collagen I [50, 51]. Integrin α1β1 also acts as a negative regulator of collagen I production and deposition, whereas integrin α2β1 promotes this process [73, 74], which is consistent with the low collagen I expression in DP cells (Figs. 1A, 1B, and 2B). This negative feedback likely contributes to maintaining the nonfibrillar, BM-like ECM within the DP niche.

### Insights into DP cell culture

Establishing effective human DP cell culture methods remains a key challenge in regenerative medicine [23, 24]. DP cells rapidly lose their ability to induce hair follicle generation in 2D culture. While BMP6 and Wnt3a treatments restore this ability in rodent DP cells [75, 76], similar approaches have not succeeded in human cells. However, 3D spheroid culture restores the ability of human DP cells to induce hair follicle generation [25, 26]. Furthermore, coculturing aggregated human DP cells with human epidermal keratinocytes has successfully generated hair follicle-like structures *in vitro* [32] and in mouse skin transplants [77], highlighting the importance of the 3D microenvironment for maintaining DP function. Given the challenges in enhancing the hair-inducing abilities of human DP cells, understanding DP-ECM interactions is critical for optimizing culture conditions. Our findings suggest that incorporating laminins containing α1, α2, and α4 chains into 3D culture could better mimic the native ECM and support DP cell function. These insights not only advance DP cell biology but also provide insights for more effective culture strategies with clinical applications in regenerative medicine.

### Experimental procedures

#### Human skin samples

Full-thickness fresh frozen human skin tissue samples were purchased from Biopredic International (Saint Grégoire, France) via KAC Co., Ltd. (Kyoto, Japan). Biopredic International complies strictly with the ethical rules for the donation and collection of human tissues according to French Law L.1245-2 CSP. Samples were isolated from the scalp, specifically the area behind the ear, of healthy Caucasian females aged 53 years after cosmetic surgery.

#### Cells and cell culture

SF-TY cells (human normal skin fibroblasts; JCRB0075) were purchased from the JCRB cell bank (Osaka, Japan) and cultured in DMEM (Sigma‒Aldrich) supplemented with 10% FBS and nonessential amino acids at 37°C, 5% CO_2,_ and 100% humidity. Primary human follicle DP cells (HFDPCs) were purchased from PromoCell (Heidelberg, Germany) and cultured in Follicle Papillae Growth Medium (PromoCell) at 37°C, 5% CO_2_ and 100% humidity. The cells were subcultured at 70–90% confluency using 0.05% trypsin/EDTA (Gibco) for SF-TY cells and a DetachKit (PromoCell) for HFDPCs. Cells at passage number 5 were used in the experiments.

#### Immunohistochemical staining

Freshly frozen human scalp skin tissues were dissected into small pieces and embedded in OCT compound within a base mold. To distinguish between different hair types, we embedded scalp skin with terminal hairs and peripheral facial skin with vellus hairs. Cryosections 100 µm thick were made using a cryostat (Leica). The skin sections were fixed with 4% paraformaldehyde (PFA)/PBS on ice for 1 h and then washed with PBS. Following washing, the skin sections were blocked with blocking buffer (10% FBS/0.1% Triton X-100/PBS) for 1 h at 4°C and then incubated with primary antibodies diluted in blocking buffer overnight at 4°C with gentle rotation (the antibody concentrations are listed in Table S1). After washing with 0.2% Tween 20/PBS for 4 h with several buffer changes, secondary antibodies and DAPI diluted in blocking buffer were added to the skin sections, which were subsequently incubated overnight at 4°C with gentle rotation. The skin sections were then washed with 0.2% Tween 20/PBS for 4 h with several buffer changes, dehydrated with 50% methanol/PBS and 100% methanol, and mounted with BABB clearing solution. Fluorescent signals were captured with a TCS SP8 confocal microscope (Leica) with a 40× oil immersion objective lens (Leica, HC PL APO 40×/1.30 Oil CS2) and LAS X software (version [BETA] 3.5.6.23723). In the scalp/facial skin samples, terminal hair follicles were located beneath the terminal hairs, and images of hair bulb regions were taken from the adipocyte layer. In contrast, in facial skin samples, vellus hair follicles were located beneath the vellus hairs, and images were taken from the dermis and upper adipocyte layer. Hair cycle stages were determined on the basis of the morphology of the hair bulb region.

#### Transmission electron microscopy

Transmission electron microscopy images of the mouse DP, which were originally obtained in our previous study [33], were reanalyzed in this study.

#### scRNA-seq data analysis

The raw sequencing data and data processing methods used were described in Yokota et al.[37]. The rds data after quality control and data integration with the list of cell population names (Gene Expression Omnibus database under accession code GSE237520) were processed with the Seurat package (version 4.1.1) in R (version 4.0.5).

To visualize the gene expression level of laminin α chains, the *DotPlot* function was used. For differential expression tests of single-cell transcriptomic data, model-based analysis of single-cell transcriptomics (MAST) [78] was used. Based on the human homologs of the mouse core matrisome genes defined in Tsutsui et al. [33, 38], we compared their expression between FB-DP and a pooled population of all dermal fibroblast subpopulations except FB-DP. Differentially expressed genes were defined as genes with adjusted *p*-values < 0.05. The volcano plot was generated using the *ggplot2* package.

To perform hierarchical clustering of the expression patterns of ECM receptor genes across dermal fibroblast populations, the average expression of each gene was calculated using the *AverageExpression* function. The clustering results were visualized as a heatmap using the *heatmap.2* function.

#### Flow cytometry analysis

The cells in subconfluent conditions were collected using a DetachKit (PromoCell) for HFDPCs and 0.05% trypsin/EDTA (Gibco) for SY-TY cells. The cells were pelleted at 200 × *g* for 5 min and then were suspended in 3% FBS/PBS at 1.0 × 10^7^ cells/ml. One million cells in 100 µl of 3% FBS/PBS were incubated with anti-human integrin antibodies at the final concentrations described in Table S1. After being washed with 3% FBS/PBS, the cells were incubated with 500 µl of 3% FBS/PBS with a 1:500 dilution of 1 mg/ml DAPI solution (Dojindo). The cells were analyzed with a SH800ZFP cell sorter (Sony). The data were analyzed with FlowJo software (BD Biosciences, version 10.9.0).

#### Preparation of extracellular matrix proteins

Human plasma fibronectin (Wako, 063-05591), human collagen I (Corning, 354243), recombinant full-length human laminins (BioLamina, laminin-111: LN111-501, laminin-211: LN211-501, laminin-332: LN332-502, laminin-411: LN411-501, laminin-511: LN511-502), and human recombinant laminin-E8 fragments (iMatrix-Palette, Matrixome, 892091) were purchased from the indicated companies.

#### Solid-phase cell adhesion assay

Solid-phase cell adhesion assays were performed as described previously [79, 80]. Briefly, 96-well cell culture plates (FALCON) were coated with 100 µl of ECM protein solution in PBS at 4°C overnight. The ECM solutions were discarded, and the plates were blocked with 1% heat-denatured BSA/PBS at 37°C for 1 h. Cells in subconfluent conditions were collected using a DetachKit (PromoCell) for HFDPCs and 0.05% trypsin/EDTA (Gibco) for SY-TY cells. The detached cells were suspended in 1% BSA/serum-free DMEM (Sigma) at a density of 6.0 × 10^5^ cells/ml. Thirty thousand cells in 50 µl of 1% BSA/serum-free DMEM were added to the ECM-coated wells and incubated for 30 min in a CO_2_ incubator at 37°C. The cells attached to the plates were fixed with 4% PFA/PBS for 20 min at room temperature and stained with 100 µl of 0.5% crystal violet (Wako) in 20% methanol for 15 min at room temperature. After washing with distilled water, the cell-bound crystal violet was extracted with 100 µl of 1% SDS for 5 min, and the OD600 was measured with an ARVO MX plate reader (PerkinElmer). For quantification, the OD600 value at 0 nM was subtracted, and the values relative to the maximum value of fibronectin were calculated. The ED50 was calculated in R (version 4.3.1) with the *drc* package [81].

For the cell adhesion-blocking experiments, integrin antibodies with cell adhesion-blocking activities (see Table S1) were added to a 6.0 × 10^5^ cells/ml cell suspension in 1% BSA/serum-free DMEM (Sigma) for 20 min at room temperature at a final concentration of 10 µg/ml. Ninety-six-well cell culture plates (FALCON) were coated with 100 µl of ECM protein solution at a concentration close to the ED50 in PBS as follows: for the experiments with HFDPCs, 1.3 nM fibronectin, 1.3 nM collagen I, 10 nM laminin-111 E8, 20 nM laminin-221 E8, 1.3 nM laminin-332 E8, 5 nM laminin-411 E8, and 0.6 nM laminin-511 E8 were used. For the experiments with SF-TY, 0.2 nM fibronectin, 0.2 nM collagen I, 5 nM laminin-111 E8, 0.6 nM laminin-332 E8, 1.3 nM laminin-411 E8, and 0.2 nM laminin-511 E8 were used. For quantification, the OD600 values were normalized to those of the No Ab condition.

#### Antibodies

The antibodies used in this study and their working dilutions are listed in Table S1.

#### Spheroid culture in extracellular matrix gels

To make cell spheroids, HFDPCs under subconfluent conditions were collected using a DetachKit (PromoCell). The detached cells were suspended in Follicle Papillae Growth Medium (PromoCell) at 3.0 × 10^3^ cells/ml. Three hundred cells in 100 µl of medium were added to a Nunclon Sphera 96-well U-bottom plate (Thermo Fisher Scientific) and incubated for 30 min in a CO_2_ incubator at 37°C. In parallel, extracellular matrix gels were prepared as described below. For the collagen I gels, 3.0 mg/ml acetic acid-solubilized collagen I derived from pig tendons (Nitta gelatin, 637-00653) was mixed with 10× MEM (Nitta gelatin) and 10× reconstruction buffer (50 mM NaOH/260 mM NaHCO_3_/200 mM HEPES, Nitta gelatin) to achieve a final concentration of 2.4 mg/ml. Mouse EHS tumor-derived Matrigel (growth factor-reduced, phenol red-free) was purchased from Corning (product number: 356231, lot numbers: 1333001, 19824002, 31123001) and used without dilution (9.2, 9.6, and 11.8 mg/ml, respectively). Fifty microliters of collagen I gel or Matrigel were added to ninety-six-well cell culture plates (FALCON) and allowed to set for 30 min in a CO_2_ incubator at 37°C. The HFDPC spheroids were transferred from 96 U-bottom plates to the surface of the gels. The spheroids were then covered with 50 µl of collagen I gel or Matrigel and incubated for 30 minutes in a CO_2_ incubator at 37°C. After incubation, the spheroids were cultured with 100 µl of Follicle Papillae Growth Medium (PromoCell) with a 1:40 dilution of HEPES (Gibco, 1 M) and a 1:100 dilution of penicillin‒streptomycin solution (Sigma, 10,000 units/ml penicillin and 10 mg/ml streptomycin) in a CO_2_ incubator at 37°C. Images were acquired via an OLYMPUS CK40 microscope with a 4× objective lens (OLYMPUS, T6-131273) and Spectman software (WRAYMER).

For the cell adhesion-blocking experiments, anti-integrin antibodies with cell adhesion-blocking activities (see Table S1) were added to the cell culture media just after spheroid formation and incubated for 20 min at room temperature. Then, the spheroids were embedded in collagen I gel or Matrigel as described above and cultured with 100 µl of Follicle Papillae Growth Medium (PromoCell) supplemented with 10 µg/ml integrin-blocking antibodies in a CO_2_ incubator at 37°C. For the spheroid invasion assays, image backgrounds were subtracted from 8-bit grayscale images using the “Subtract Background” function in Fiji/ImageJ software (ver. 2.0.0-rc-66), and the spheroid surface was enhanced via the “Enhance contrast” function. The spheroid areas were then manually identified using polygon tools.

#### Statistics and reproducibility

No statistical method was used to predetermine the sample size. All the statistical analyses were performed using R (version 4.3.1), and plots were generated using the *ggplot2* package (version 3.4.3). Statistically significant differences are indicated in each figure (\**p*<0.05, \*\**p*<0.01, and \*\*\**p*<0.001). We show representative micrographs that were obtained from at least two biological replicates except for Fig. S1B.

## Data availability

All image data in this study have been stored in the SSBD repository at https://doi.org/10.24631/ssbd.repos.2025.01.415. The scRNA-seq data in this study is available at Gene Expression Omnibus database under accession code GSE237520. The source data for all figures are available at the Mendeley Data under the DOI: 10.17632/28x5xjf8f9.1. All data that support the findings of this study are available within the paper and its supplementary information files.

## Acknowledgments

We thank members of the Fujiwara laboratory for valuable discussions and reagents. We also thank Kiyotoshi Sekiguchi and Yukimasa Taniguchi for their technical advice. This work was supported by RIKEN intramural funding, the BDR Organoid Project, the JSPS KAKENHI (19K22631, 20H03706, 22K19453, 25H01053), the MEXT KAKENHI (23H04927, 23H04928), and the JST CREST (JPMJCR1926) (all to H. Fujiwara). H. Machida was supported by the RIKEN Junior Research Associate (JRA) and a JSPS Research Fellow (22KJ2160).

## Author contributions

H. Machida designed and carried out the experiments, analyzed the data, and wrote the initial draft. J. Yokota generated the scRNA-seq data and supported the data analysis. H. Fujiwara conceived the study, designed and supervised the study, and edited the initial draft.

## Declaration of interests

H. Fujiwara and H. Machida are inventors on a patent application concerning the *in vitro* culture of dermal papilla cells and related cell aggregates. The other authors declare no competing interests.

## Abbreviations

DP: dermal papilla
HG: hair germ

## Notes

https://doi.org/10.24631/ssbd.repos.2025.01.415

https://www.ncbi.nlm.nih.gov/geo/query/acc.cgi?acc=GSE237520

