## Supplemental Materials for "Basement membrane components define the microenvironment of aggregated fibroblasts in the skin and support their aggregation *in vitro*"

**Supplementary Data**

A

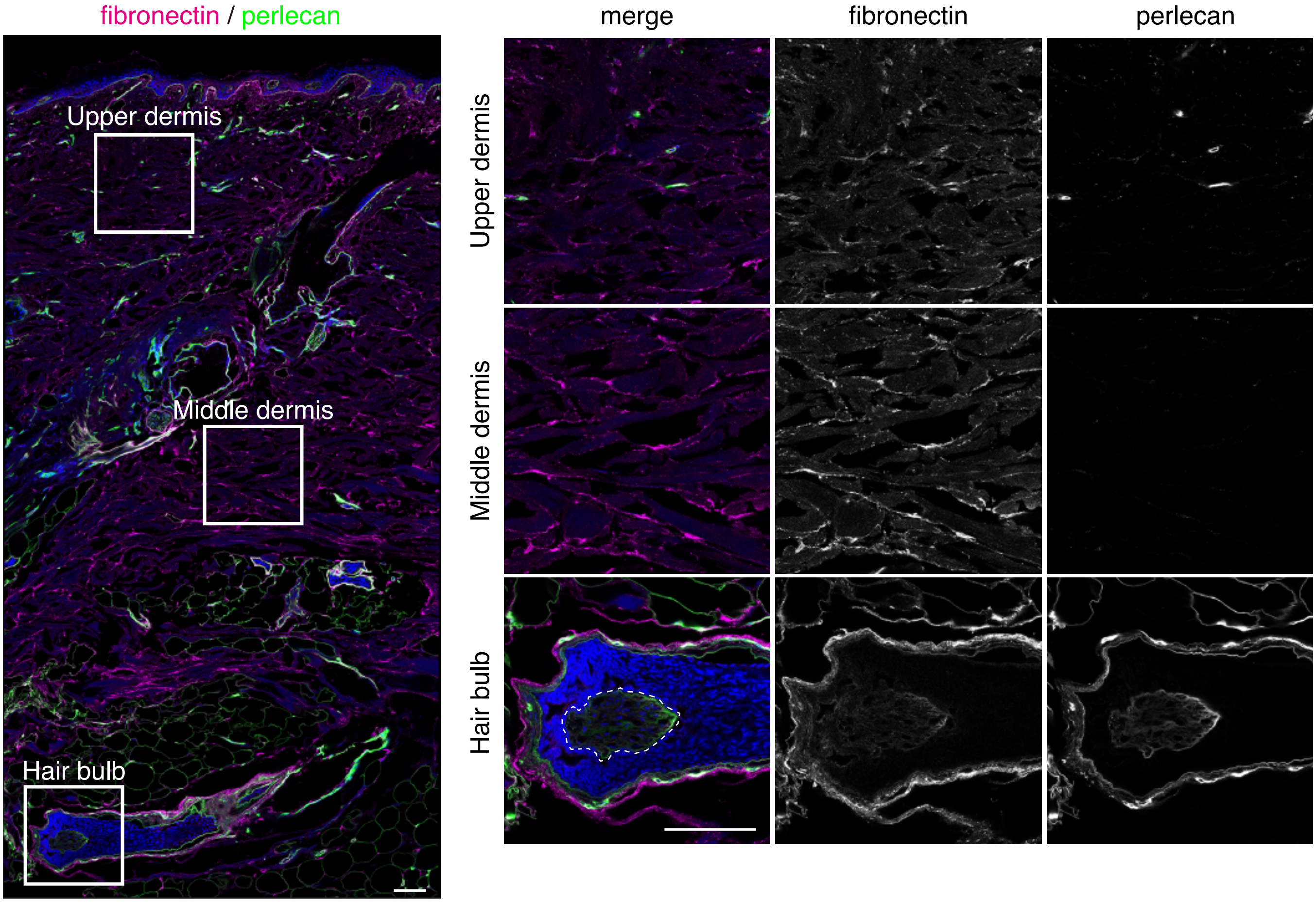
fibronectin / perlecan
Upper dermis
Middle dermis
Hair bulb
merge
fibronectin
perlecan
Upper dermis
Middle dermis
Hair bulb

B

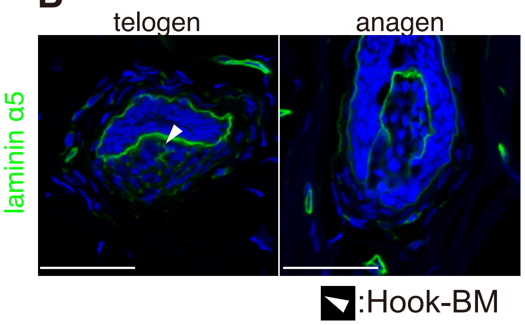
telogen
anagen
laminin α5
:Hook-BM

Fig. S1

**Fig. S1. Tissue distribution of ECM proteins in the human DP.**

(A) Immunohistochemical staining of the interstitial fibrillar ECM component fibronectin (magenta) and the ubiquitous BM protein perlecan (green) in human female scalp skin with terminal hair follicles. Nuclei are stained with DAPI (blue). The white squares indicate the magnified regions shown on the right. In the hair bulb panel, the dashed lines outline the DP. Scale bar, 100 µm. (B) Immunohistochemical staining of laminin α5 (green) in the hair bulb region of human female facial vellus hair follicles. The arrowhead indicates a hook-like distribution of laminin α5 in the DP. Scale bar, 50 µm.

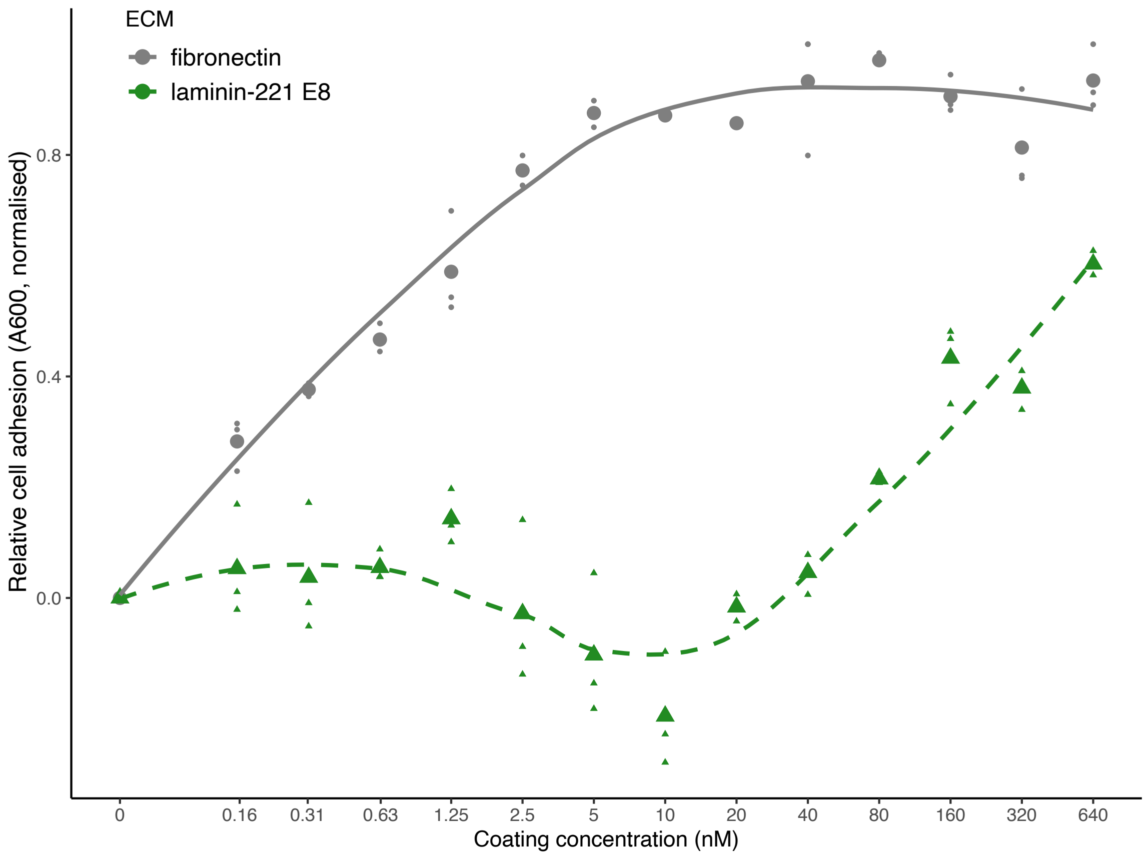
ECM
fibronectin
laminin-221 E8
Relative cell adhesion (A600, normalised)
0.8
0.4
0.0
0
0.16
0.31
0.63
1.25
2.5
5
10
20
40
80
160
320
640
Coating concentration (nM)

Fig. S2

**Fig. S2. The adhesion of SF-TY cells to laminin-221 E8 does not reach a plateau even at 640 nM.**

Dose–response curves showing the relative cell adhesion in solid-phase cell adhesion assays using the human normal skin fibroblast line SF-TY. The absorbance at 600 nm represents the relative number of adhered cells. Smaller dots indicate normalized values of adhered cells relative to the maximum effect observed on fibronectin for each technical replicate. Larger dots indicate the mean values for each ECM protein, and the colored lines show their corresponding fitting curves.

**Table S1: Antibodies used in this study**

| Designation | Source | Identifiers | Application<br>Working dilution |
|---|---|---|---|
| Anti-collagen type I<br>(clone EPR7785) | Abcam | Ab215969 | IHC (1:1000 [final conc. 1.0 μg/ml]) |
| Anti-collagen type IV<br>(clone PHM-12, CIV22) | Thermo | MA5-14100 | IHC (1:200) |
| Anti-fibronectin | Abcam | Ab2413 | IHC (1:200) |
| Anti-integrin alpha1<br>(clone CL7207) | Abcam | Ab243032 | IHC (1:500 [2.0 μg/ml]) |
| Anti-integrin alpha1-PE<br>(clone TS2/7) | eBioscience | 12-9490-41 | FACS (1:20 [5.0 μg/ml]) |
| Anti-integrin alpha1<br>(clone FB12) | Merk | MAB1973Z | Blocking (1:100 [10.0 μg/ml]) |
| Anti-integrin alpha2<br>(clone P1E6) | DSHB | | IHC (1:160 [6.3 μg/ml])<br>Blocking (1:100 [10.0 μg/ml]) |
| Anti-integrin alpha2-PE<br>(clone P1E6-C5) | BioLegend | 359307 | FACS (1:20 [5.0 μg/ml]) |
| Anti-integrin alpha3<br>(clone P1B5) | DSHB | | IHC (1:400 [2.5 μg/ml])<br>Blocking (1:100 [10.0 μg/ml]) |
| Anti-integrin alpha3-APC<br>(clone P1B5) | eBioscience | 17-0494-42 | FACS (1:20 [1.3 μg/ml]) |
| Anti-integrin alpha5<br>(clone P1D6) | DSHB | | IHC (1:500 [2.0 μg/ml])<br>Blocking (1:100 [10.0 μg/ml]) |
| Anti-integrin alpha5-PE<br>(clone IIA1) | BD bioscience | 555617 | FACS (1:5 [5.0 μg/ml]) |
| Anti-integrin alpha6<br>(clone GoH3) | BioLegend | 313637 | IHC (1:200 [5.0 μg/ml])<br>Blocking (1:100 [10.0 μg/ml]) |
| Anti-integrin alpha6-PE/Cy7<br>(clone GoH3) | BioLegend | 313621 | FACS (1:20 [5.0 μg/ml]) |

| | | | |
|---|---|---|---|
| Anti-integrin alpha7 (clone E-2) | SantaCruz | Sc-515716 | IHC (1:200 [1.0 μg/ml]) |
| Anti-integrin alpha7-PE (clone 3C12) | MBL | K0046-5 | FACS (1:1.25 [4.4 μg/ml]) |
| Anti-integrin alpha7 (clone 9.1 ITGA7) | DSHB | | Blocking (1:100 [10.0 μg/ml]) |
| Anti-integrin alpha8 | R&D | AF4076 | IHC (1:500 [0.4 μg/ml]) |
| Anti-integrin alpha9 | R&D | AF3827 | IHC (1:160 [6.3 μg/ml]) |
| Anti-integrin alpha9-PE (clone Y9A2) | BioLegend | 351605 | FACS (1:20 [10.0 μg/ml]) |
| Anti-integrin alphaV | R&D | AF1219 | IHC (1:80 [5.0 μg/ml]) |
| Anti-integrin alphaV-PE (clone NKI-M9) | BioLegend | 327909 | FACS (1:20 [2.5 μg/ml]) |
| Anti-integrin alphaV (clone L230) | DSHB | | Blocking (1:100 [10.0 μg/ml]) |
| Anti-integrin beta1 (clone P5D2) | DSHB | | IHC (1:1000 [1.0 μg/ml]) Blocking (1:100 [10.0 μg/ml]) |
| Anti-integrin beta1-PE (clone MAR4) | BD bioscience | 561795 | FACS (1:5 [16.7 μg/ml]) |
| Anti-integrin beta4 | Sigma | HPA036348 | IHC (1:1000 [1.0 μg/ml]) |
| Anti-laminin alpha1 (clone G-12) | SantaCruz | Sc-74418 | IHC (1:400 [0.5 μg/ml]) |
| Anti-laminin alpha2 (clone 5H2) | Merk | MAB1922 | IHC (1:1000) |
| Anti-laminin332 | Abcam | Ab14509 | IHC (1:400 [2.5 μg/ml]) |
| Anti-laminin alpha4 (clone 3H2) | Abnova | MAB7869 | IHC (1:200 [0.5 μg/ml]) |

| | | | |
|---|---|---|---|
| Anti-laminin alpha5 (clone 4C7) | Abcam | Ab17107 | IHC (1:10 [200 µg/ml]) |
| Anti-perlecan (clone A7L6) | Millipore | MAB1948P | IHC (1:500 [2.0 µg/ml]) |
| Goat anti-mouse IgG2a, Alexa555-conjugated | Life Technologies | A21127 | IHC (1:1000) |
| Goat anti-mouse IgG1, Alexa647-conjugated | Life Technologies | A21241 | IHC (1:1000) |
| Goat anti-rat IgG, Alexa555-conjugated | Life Technologies | A21434 | IHC (1:1000) |
| Goat anti-rabbit IgG, Alexa647-conjugated | Life Technologies | A21245 | IHC (1:1000) |
| Donkey anti-goat IgG, Alexa555-conjugated | Life Technologies | A21432 | IHC (1:1000) |
| Donkey anti-rabbit IgG, Alexa647-conjugated | Life Technologies | A31573 | IHC (1:1000) |
| Mouse IgG1 isotype control-PE (clone MOPC21) | BD bioscience | 555749 | FACS (1:5 [1.0 µg/ml]) |
| Mouse IgG1 isotype control-APC (clone MOPC21) | BioLegend | 400121 | FACS (1:160 [1.3 µg/ml]) |
| Rat IgG2a isotype control-PE/Cy7 (clone RTK2758) | BioLegend | 400521 | FACS (1:40 [5.0 µg/ml]) |
| Mouse IgG2a Isotype control-PE (clone MOPC173) | BioLegend | 400213 | FACS (1:80 [2.5 µg/ml]) |
| Mouse IgG isotype control (clone P3.6.2.8.1) | eBioscience | 16-4714 | Blocking (1:100 [10.0 µg/ml]) |
